# Insula cortex gates the interplay of action observation and preparation for controlled imitation

**DOI:** 10.1101/2020.11.05.370718

**Authors:** Megan E. J. Campbell, Vinh T. Nguyen, Ross Cunnington, Michael Breakspear

## Abstract

Perceiving, anticipating and responding to the actions of another person are fundamentally entwined processes such that seeing another’s movement can prompt automatic imitation, as in social mimicry and contagious yawning. Yet the direct-matching of others’ movements is not always appropriate, so this tendency must be controlled. This necessitates the hierarchical integration of the systems for action mirroring with domain-general control networks. Here we use functional magnetic resonance imaging (fMRI) and computational modelling to examine the top-down and context-dependent modulation of mirror representations and their influence on motor planning. Participants performed actions that either intentionally or incidentally imitated, or counter-imitated, an observed action. Analyses of these fMRI data revealed a region in the mid-occipital gyrus (MOG) where activity differed between imitation versus counter-imitation in a manner that depended on whether this was intentional or incidental. To identify broader cortical network mechanisms underlying this interaction between intention and imitativeness, we used dynamic causal modelling to pose specific hypotheses which embody assumptions about inter-areal interactions and contextual modulations. These models each incorporated four regions - medial temporal V5 (early motion perception), MOG (action-observation), supplementary motor area (action planning), and anterior insula (executive control) – but differ in their interactions and hierarchical structure. The best model of our data afforded a crucial role for the anterior insula, gating the interaction of supplementary motor area and MOG activity. This provides a novel brain network-based account of task-dependent control over the integration of motor planning and mirror systems, with mirror responses suppressed for intentional counter-imitation.

## 1 Introduction

Interacting with others is the quintessential human experience. Dyadic interactions rely upon anticipating, perceiving and interpreting the actions of another person while concurrently preparing and executing one’s own actions. The understanding of perception-action integration has been heavily influenced by research on so-called ‘mirror neurons’, first documented in non-human primates (Ferrari and Rizzolatti, 2014). The human “mirror neuron system” is a constellation of fronto-parietal regions that were traditionally thought to subserve motor control yet have since been shown to also respond to observing the actions of others (Molenberghs et al., 2012), and forms part of a wider action observation network which extends into the temporo-occipital cortices (Caspers et al., 2010). Previous research suggests that the mapping of an external agent’s actions on to the sensorimotor system helps pre-empt the likely movements, affordances, and goals of that agent (Ferrari and Rizzolatti, 2014; Rizzolatti and Sinigaglia, 2010). Furthermore, this “motor resonance” prompts self-made actions in response to the actions of an external agent, particularly priming imitative and complementary movements (Molenberghs et al., 2009). Automatic imitation is thought to reflect this internal mirroring of visuo-motor stimuli that either interferes or enhances response preparation (Heyes, 2011). Yet humans do not simply copy each other. Rather, contextual cues to the relevance of mirroring another’s actions for particular goals influence response preparation (e.g. a boxing match distinctly requires complementary and even opposing responses). The preparatory inhibition of automatic imitation in contexts where copying another’s movement is counter-productive has become the topic of substantial research (Bardi et al., 2015; Brass et al., 2009; 2001; Cross and Iacoboni, 2014a; 2014b; Cross et al., 2013).

The mirror neuron system is integrated within a hierarchy of other functional systems, including executive control networks (Campbell and Cunnington, 2017). Such integration must be adaptive, balancing automatic mirroring versus inhibition in a dynamic and contextually sensitive manner. We recently used functional magnetic resonance imaging (fMRI) to disambiguate mechanisms for the top-down suppression of mirror representations when imitation was contrary to task demands (*intentional* counter-imitation), as distinct from a response that *incidentally* mismatched a task-irrelevant stimulus (Campbell et al., 2018). The paradigm was thus a modified automatic imitation, stimulus-response (SR) compatibility paradigm (Brass, Bekkering, Wohlschläger, & Prinz, 2000; Cross & Iacoboni, 2014b). The classical SR compatibility effect is observed behaviourally as faster reaction times when performing a response that is congruent with the imperative stimulus, than when the response is incongruent, even if this SR congruence is task-irrelevant (Prinz, 1997; Zwickel and Prinz, 2012). When applied to action mirroring and automatic imitation this reaction time benefit occurs for imitatively congruent SR pairs, over non-imitative/incongruent SR pairs (Brass, Bekkering, Wohlschläger, & Prinz, 2000; Heyes, 2011). Our analysis of this fMRI dataset revealed an assemblage of frontal regions that were recruited more strongly when counter-imitation was intentional than when it was incidental. These regions derived largely from the “cingulo-operculum control network” (Dosenbach et al., 2008). Furthermore, a cluster of regions in the middle occipital gyrus and angular gyri displayed an interaction between task-relevance (intentional versus incidental) and counter-imitation versus imitation. We interpreted this interaction as reflecting the boosting of mirror-matched visual representations of observed actions when these were explicitly task relevant, to aid the preparation of an imitative response to the stimulus action (Campbell et al., 2018). This is consistent with the contribution of the inferior parietal lobule to the frontoparietal mirror neuron system for visuo-motor integration of observed actions (Ferrari and Rizzolatti, 2014; Rizzolatti and Sinigaglia, 2010), yet also extends beyond the classical ‘mirror neuron’ front-parietal areas and into the wider action observation network (Caspers et al., 2010; Molenberghs et al., 2012).

Whilst informative, task-related activations do not reveal the broader network interactions enacting the sensitive and contextual nature of adaptive mirror responsivity. These rely on directed and mutual interactions, and their contextual up- or down-regulation. Here we used dynamic causal modelling (DCM) for fMRI to map out a biologically plausible neural model to understand adaptive and context-dependant mirroring by exploring the cortical mechanisms underlying our previous analysis (Campbell et al., 2018).

## 2 Methods

### 2.1 Participants and behavioural paradigm

Twenty-four neurologically healthy, right-handed adults (13 females; mean age = 23.5, SD=3.3 years) participated in this study (Campbell et al., 2018). The study had Institutional human research ethics approval and participants provided written informed consent. Behavioural response recordings were unreliable for four participants. As such, behavioural effects were tested on a subsample of 20 participants. However, the fMRI analysis was based on the full sample of 24.

As previously described, this task required participants to respond to action observation with concurrent action execution (Campbell et al., 2018). Stimulus and response actions were based on two contradictory movements: hand opening or hand closing gestures (Fig. 1A). Participants were instructed to respond in one of four ways – to open or close their own hand, or to copy or oppose the observed action. This congruence (match/mismatch) between observed and executed actions, was controlled to be equally likely on any trial. Distinct from other SR compatibility paradigms (Brass, Bekkering, Wohlschläger, & Prinz, 2000; Cross & Iacoboni, 2014b), we manipulated the task-relevance of stimulus-response (SR) congruence, with automatic (and counter-imitation) or *incidental imitation* compared to *intentional imitation* (and counter-imitation). Our paradigm was designed to distinguish preparatory versus reactive inhibition of mirroring by separating the processes of *incidentally* observing a mismatching action from *intentionally* counter imitating it. As we reported previously (Campbell et al., 2018), we showed a classic behavioural effect of slowed reaction times for mismatched versus matched responses, across both incidental and intentional conditions (Fig. 1B).

**Figure 1:**
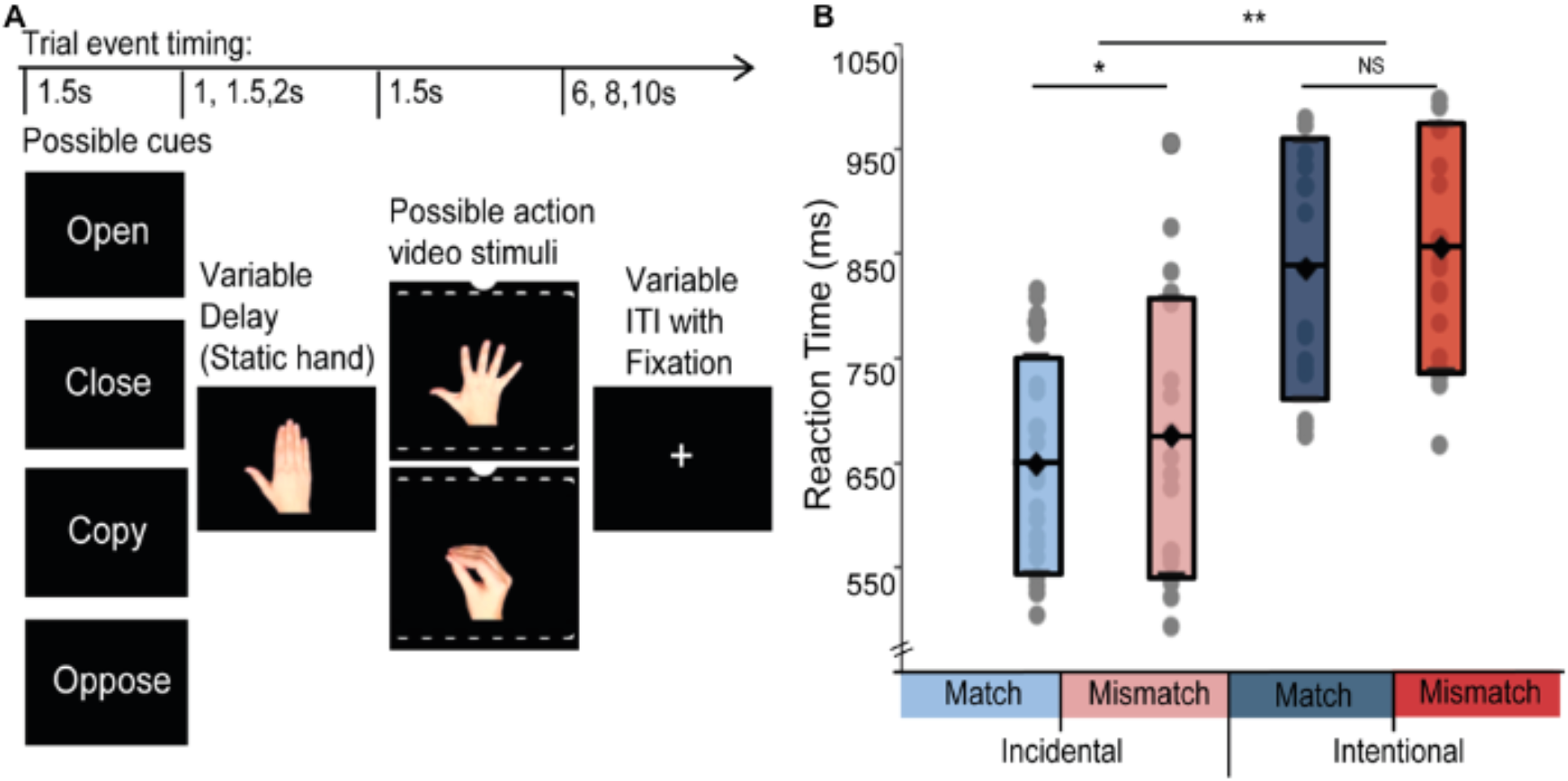
**A. Event related experimental design.** Comprised four trial types in a 2×2 factorial design: task-relevance (incidental/intentional) x SR congruence (imitation/counter-imitation). Participants were instructed to perform the cued response as soon as the movement in the video was detected. Thus SR pairs being the same or opposing actions was either *incidental* (when participants were cued to perform ‘open’ or ‘close’ movements regardless of the presented action) or *intentional* (with the stimulus action being explicitly task-relevant given participants were cued to either ‘copy’ or ‘oppose’ the presented action). ITI: inter-trial interval. **B. Reaction time effects:** Box plots indicate group mean and standard deviation, grey dots showing individual participant means for each condition. ** p<0.001; *p<0.05; NS: Non-significant. Note: reaction time data analysed for a subsample, n=20, as response recording was unreliable for 4 participants.

### 2.2 Data-acquisition

Functional MRI images were acquired with gradient-echo echo-planar imaging simultaneous multi-slice acquisition sequence with the following parameters: 44 axial slices (multiband acceleration factor 4); echo-time (TE)=32.0ms; repetition-time (TR)=700ms; flip-angle (FA)=70°; pixel bandwidth 1698; field-of-view=200×200mm; 74×74 voxel matrix; 3.0×3.0×3.0 mm3 with 10% slice gap, whole brain coverage. Structural scans were acquired for alignment and coregistration to MNI space (MP2RAGE sequence, TE=2.32ms, TR=1900ms, FA=9°, 256×256 cubic matrix, voxels = 0.9×.09×0.9mm^3^).

Infrared motion-capture was used to record reaction time and monitor accuracy (Qualisys Motion Capture system with Qualisys Track Manager software). Two wall-mounted infrared cameras recorded the changing position of markers attached to participants’ thumb and index finger (250 frames per second) from 200ms before stimulus presentations until the end of the response period. Thus the X, Y, Z trajectories of the markers (as moving X, Y, Z plane coordinates) indicated action onsets relative to stimulus-onset and was used to quantify reaction-times. The movement onset for each participant, for each trial was defined as the time point when acceleration exceeded one-tenth of a standard deviation of that participant’s moving average (Campbell et al., 2018; Mehrkanoon et al., 2014) reflecting a marked deviation from the participant’s starting rest position. The motion capture recordings from four participants were unreliable with less than a 30 percent of trials in each condition, and so reaction-time analysis was based on a subsample of 20 participants.

### 2.3 fMRI Pre-processing and General Linear Modelling

The functional MRI were acquired and pre-processed as previously described (Campbell et al., 2018). Pre-processing included: spatial realignment (6-degree affine transformation to the first image of the first scan for head-movement correction); co-registration of each individual’s mean functional MRI image to their T1 image; spatial normalisation based on T1 image (nonlinear transformation to MNI space using the SPM8 segment process); re-slicing to 2 x 2 x 2 mm^3^, and spatial smoothing (6mm FWHM isotropic Gaussian kernel); all applied in SPM8 (Wellcome Department of Imaging Neuroscience, Institute of Neurology, London, www.fil.ion.ucl.ac.uk/spm) and implemented in MATLAB (R2015b, Mathworks).

The GLM design matrix was built on the four task conditions: *Intentional Imitation, Incidental Imitation, Intentional Counter-imitation* and *Incidental Counter-imitation* (Campbell et al., 2018), Each condition was compared to the implicit baseline of fixation during intervals between trials. Haemodynamic response functions were modelled for the onset of action stimuli, covering periods of both action preparation and response execution. Here we used a 2×2 Flexible Factorial design to test for the separate main effects of SR congruence (imitation/congruent versus counter-imitation/incongruent) and of task-relevance (intentional versus incidental) and the interaction between these factors.

### 2.4 Modelling Effective Connectivity

We used dynamic causal modelling (DCM Friston et al., 2003a) to examine effectivity connectivity between the brain areas implicated in task-dependent modulation of mirror representations within the action observation network. DCM embodies effective connectivity by causal terms in a differential equation (Friston et al., 2003b; Stephan & Friston, 2010) with one state for each node and user specified parameters for: i) the driving inputs onto specific regions, ii) intrinsic connections between regions, and iii) task-dependent modulators of these interactions in order to frame plausible hypotheses for cortical networks (Stephan et al., 2008; Kahan and Foltynie, 2013; Stephan et al., 2010). Nonlinear DCM incorporates hierarchical relationships between regions, such that one region influences the effective connections between another pair of regions (Stephan et al., 2008). Given our goal of explaining an interaction effect between SR congruence and intentionality, we focused on positing nonlinear models of modulation, with only one bilinear design for comparison.

DCM was implemented with SPM12 in MATLAB 2015b, and involves two steps. Firstly, a user-specified model design based on task-related BOLD responses reflected in GLM analysis, combined plausible network hypotheses motivated by prior knowledge. Finally, data-driven model comparison based on the probabilistic evidence for each model within this user-defined model space to identify the model which best explains the observed data (Stephan, et al. 2010).

#### 2.4.1 Model specification

Firstly, BOLD-signal time courses were extracted from voxels-of-interest (VOIs) that were selected based on our group-level flexible factorial contrast as showing significant task-related responses to the main and interaction effects of our paradigm (See Results Fig. 2, and Supplementary Information section S1.1 for details of node selection). These VOIs were functionally defined for each participant, as 6mm spheres around these group peak-coordinates within structurally defined areas as per Automatic Anatomical Labelling toolbox, AAL (Tzourio-Mazoyer et al., 2002). Our network nodes were restricted to left-hemisphere areas, being contralateral to the hand used to perform the task.

**Figure 2:**
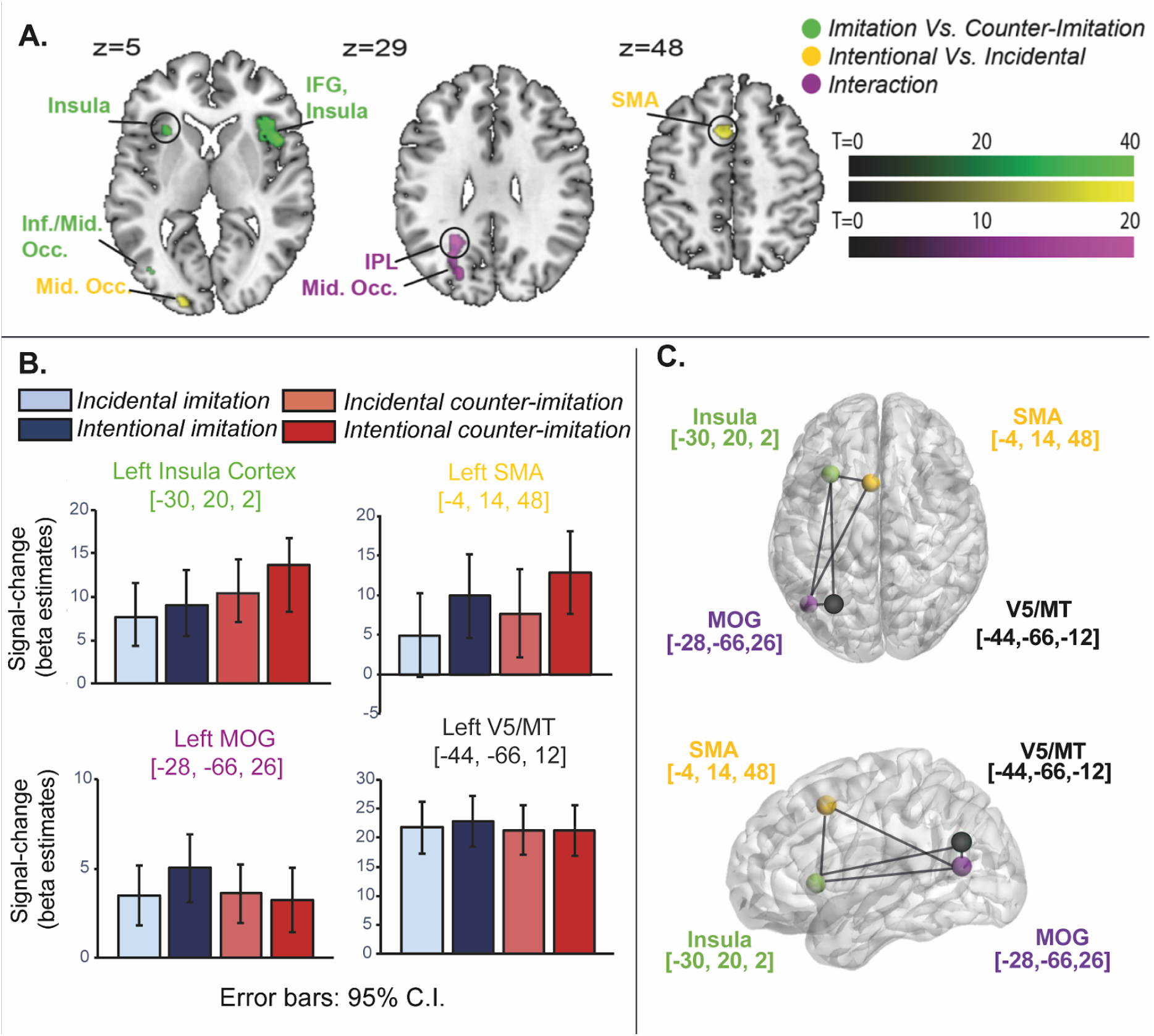
BOLD task effects. A) Main effects of SR congruence (match/mismatch) and task-relevance (intentional/incidental; p<0.001, FWE-corrected) and the interaction between these factors (p<0.05, FWE-corrected); all with cluster extent threshold of 25 voxels. B) Beta-estimates by task conditions. Colours denote conditions, blues: imitation, reds: counter-imitation; lighter shades: incidental responses; darker shades: intentional responses. All regions responded to the task itself, with the V5 peak voxel showing equal change across conditions. The left insula cortex peak [−30, 20, 2] displayed a main effect of SR congruence, and was greatest for SR mismatch. The SMA peak [−4, 14, 48] BOLD showed a main effect of task-relevance, with greatest change for intentional over incidental. The MOG peak [−28, −66, 26] showed an interaction between these two factors, with strongest effect evoked for intentional imitation, and a weaker effect for all other conditions. C) Dynamic causal models were constructed to explain these effects building on a basic model network architecture between these four regions (image created in Brainnet Viewer, Xia, Wang, & He, 2013).

Our final network design, on which all our subsequent models of effectivity connectivity are built, is hence composed of the following nodes: Area V5/MT, MOG, SMA and anterior insula cortex from the left hemisphere, with respective peak coordinates: [−44 −66, 12]; [−28, −66, 26]; [−4, 14, 48]; [−30, 20, 2] in MNI space (refer to Fig.2 and Supplementary Information Tables S1,2 for peak statistics). Our hypotheses were framed in terms of four integrated systems, with a node from each functional network captured by these VOIs of our base-model: 1. low-level visual perception of motion (V5/MT); 2. action observation processing (MOG); 3. motor preparation (SMA); and 4. higher-order cognitive control functions (anterior insula).

Networks of effective connectivity between these nodes and their modulation were specified to explain our GLM results in terms of functional integration. These are motivated and described below, in Results, Section 3.2 (further details for the background rationale and the literature base for these choices are provided in Supplementary Information S1).

#### 2.4.2 Model Selection

We performed model-wise Bayesian Model Selection (BMS) using random-effects analysis (Stephan et al., 2009; Stephan & Friston, 2010) to select from our 48 hypothesised DCMs, the model that was most likely to explain the task-dependent modulation of mirror responses to observed actions in our data. We employed random effects analysis, which assumes participants may have significant neural or strategic differences to allow for maximal inter-subject variability (Stephan et al., 2010; Marreiros et al., 2010; Penny, 2012).

## 3 Results

As previously reported (Campbell et al., 2018), our behavioural dataset revealed stimulus-response congruence effects in reaction times (Fig. 1B). Responses were faster for matching SR pairs than for mismatching (F_(1,19)_ = 5.817, *p*=0.026, partial *η*2 =0.234) and faster for incidental compared to intentional responses (F_(1,19)_, *p*< .0001, partial *η*2 = 0.885). There as no significant interaction between intention/incidental and match/mismatch task factors (F_(1,19)_ =0.128, *p*=0.724).

Factorial GLM analyses of the functional MRI data have been reported previously (Campbell et al., 2018) but are repeated here since the functionally segregated responses form the bases for our DCM models. Significant differences were observed between imitative and counter-imitative actions bilaterally in the anterior insula and left inferior frontal gyrus, and between intentional and incidental responses in the supplementary motor area (SMA; Fig. 2). Activations in these frontal regions were greater for intentional counter-imitation than for incidental mismatching (Fig. 2B). There was an interaction effect for SR congruence and task-relevance within a cluster across angular gyrus of the inferior parietal lobule and MOG (purple cluster, Fig. 2A). The local maxima of significant signal-change within this parietal-occipital cluster was in the MOG (−28, −66, 26), nearing the intraparietal sulcus (Choi et al., 2006). Summary of peak statistics for main effects and interactions are detailed in the Supplementary Information Tables S1 and S2.

### 3.1 Dynamic causal modelling

To study the mechanisms underlying these effects, we constructed 48 distinct dynamic causal models (DCMs) comprised of four families, each of which embodied specific hypotheses accounting for SR congruence and task-relevance, and the interaction between them. Network nodes were selected from four regions: medial temporal area V5, MOG, SMA and anterior insula. We restricted our model to left-hemisphere areas, being contralateral to the hand participants used to perform the task. The edges between the four nodes of our model were identical across all models in the model-space (see Supplementary Information section S1.2 for further details). The MOG and insula regions were modelled as being fully connected, with bidirectional connections linking both regions to all other regions (Fig. 2C). To capture the distinct phases of early action observation and motor planning, edges between the SMA and V5 were not included. Thereafter all families modelled cleaved into the following families:

#### A: Driving inputs

The driving inputs to the network (the imperative stimulus of the observed movement onset) framed two hypotheses testing whether a single-input or dual-inputs model had most explanatory power. The first half of model space (Fig. 3A1, DCMs 1 to 24) included both a driving input to the V5 node, and a driving-input to the SMA capturing lower level motor plans. The second half of the model space (Fig. 3A2, DCMs 35 to 48) included only a visual driving input to area V5.

**Figure 3:**
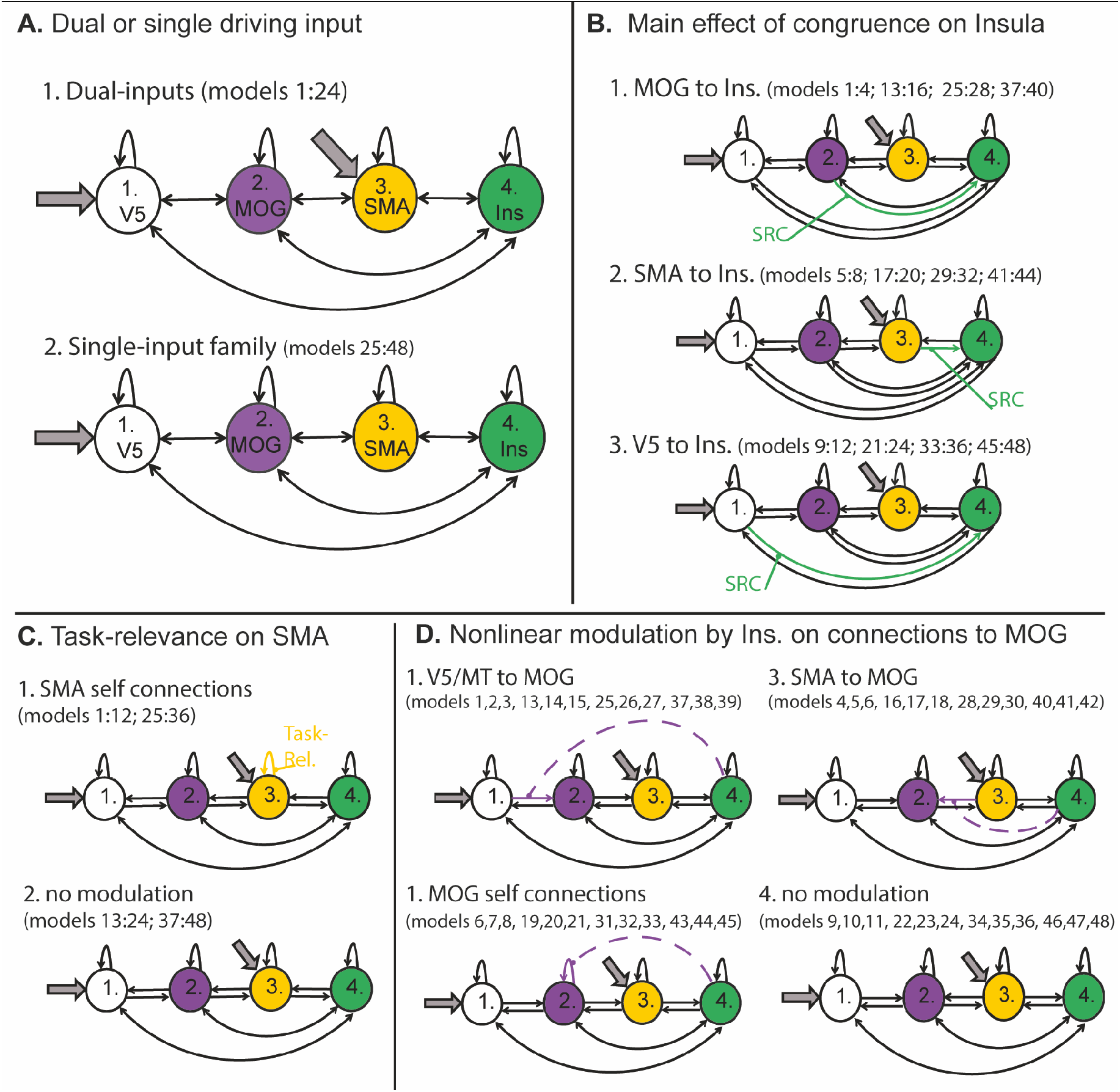
Model space of 48 DCMs constructed with each combination of separate hypotheses for: A) 2 hypotheses for driving inputs with either single (to V5) or dual (to both V5 and SMA) designs (grey arrows indicate driving inputs); B) 3 designs testing for SR congruence effect modulating connections to the insula cortex; C) 2 designs testing for the modulation of the SMA self-connection by task-relevance (intentional/incidental responses); and D) 4 designs to test for the insula gating incoming connection to the MOG, via nonlinear modulation (purple dashed lines indicate gating connection, absence indicates no modulation).

#### B. Main effect of SR congruence on the Insula

The next family of models embodied different hypotheses to account for the greater activity in the insula during SR congruence. This was achieved through parametric modulation of the connections to the insula according to the presence of SR congruence: Separate model families included modulation of either the connection of the SMA to the insula (Fig. 3.B1), the MOG to insula (B2) or the V5 to insula connection (B3). Thus, this family subdivision tested whether SR congruence was processed within the insula in terms of low-level motion perception (V5 to Ins, B1), motor preparation (SMA to Ins., B2) or action observation processes (MOG to Ins., B3).

#### C: Task-relevance on SMA

The conditions of intentional/incidental responses evoked a significantly increased BOLD response in the SMA (see GLM results in Section 3: Fig. 3.A, B). To model the mechanisms underlying this effect, we tested whether the intrinsic (self-connections) within the SMA were modulated by task-relevance, or not (Fig. 3C).

#### D. Interaction of SR congruence and task-relevance within the MOG

Nonlinear effects can be accommodated within DCM (Stephan et al., 2008, see Methods Section 2.4 for detail) to allow for internally generated interaction effects (Fig. 3D). Here, non-linear connections were introduced to account for the interaction effect within the MOG region of interest. This allows for testing how the insula cortex may exert top-down modulation over visual representations of actions. Four alternative hypotheses were incorporated: the insula gating the V5 to MOG connection (low level visual input to the action observation network, D1); or the insula gated the SMA to MOG connection (integration of motor planning action observation, D2); or the insula gating the intrinsic self-connections within the MOG node (modulating activity within this region, D3); or, a null effect lacking non-linear modulation (D4).

All combinations of these various hypothesis yielded the 48 distinct models tested by comparing the likelihood of all models as estimated by variational Bayes using Free Energy to implement Bayesian Model Selection (BMS).

The most likely model of effective connectivity for our data was the fifth model specified, with an exceedance probability of 0.190, and a protected exceedance probability of 0.348 (Fig. 4). The model combines the following specific features of each family: Driving inputs to both V5 and SMA (model-space design Fig. 3, A1); modulation of stimulus congruency on the connection from the SMA to the insula (B2); modulation of task-relevance (intentionality) on the self-connection of the SMA (C1); and crucially – to explain the interaction between SR and intentionality – a nonlinear gating of the connection from SMA to MOG, by the insula (D2).

**Figure 4:**
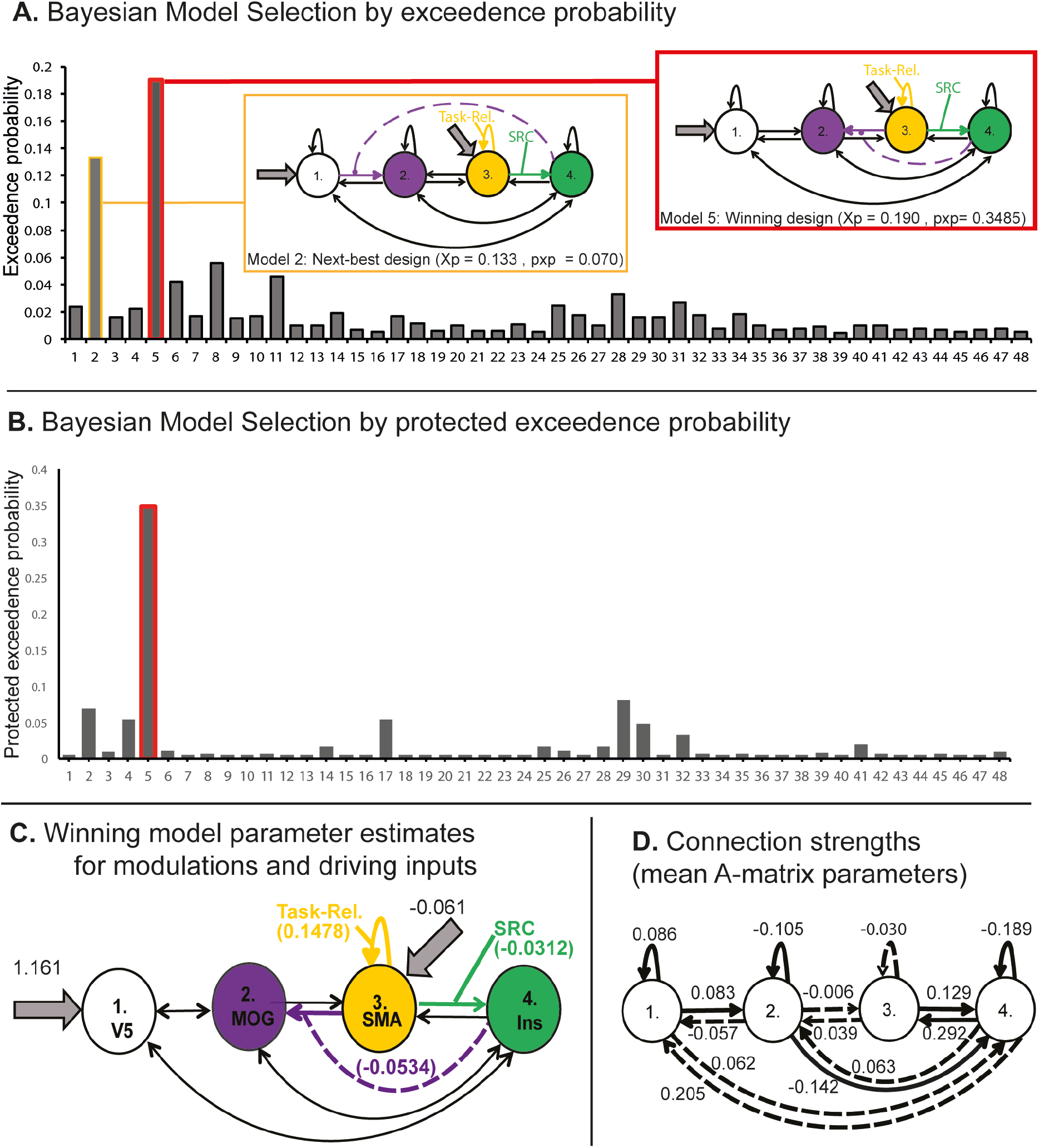
DCM model inversion. A) BMS exceedance probabilities of all 48 models. B) BMS protected exceedance probabilities. Bayes Omnibus Risk (BOR) = 0.15. Break-out boxes: Red box is the wining model (number 5); yellow box is the next-best performing model (number 2). C) Mean parameter estimates for task modulations and nonlinear effects (B and D matrices, respectively), and mean parameter estimates for driving inputs (C-matrix); D) Mean parameter estimates (Ep) for intrinsic connections (A-matrix parameters), solid lines indicating connections strengths of significant difference from zero (p<0.05).

The model parameter estimates embody the mechanisms underlying task-related effects (Fig. 4C,D). To recap, the response of the MOG, taken to reflect action observation processing, was its greatest during intentional imitation, i.e. the only task condition for which a mirror-matched response was explicitly task relevant. The encoding of the intentional/incidental SR mapping (Fig.3) is reflected in the stronger (additive) contribution to its self-connection by the SR modulatory effect, which in essence increases the BOLD response when the SR mapping is intentional, not incidental.

The effective connection from the SMA to the MOG represents the integration of the visual representation of the observed action and its task-relevance to motor preparation. The insula response reflected stimulus-response mismatch. So, by gating the SMA to MOG connection through a negative nonlinear effect (Fig. 4C), the insula suppresses the interplay between action observation and motor-preparation, when there is both a stimulus-response conflict detected (by the insula) and when the SMA is preparing a stimulus-relative response (intentional rather than incidental).

## 4 Discussion

Dyadic and higher order human interactions rest upon the conjoint observation and planning of action, with particular importance paid to the context-specific imitation and counter-imitation of others. We sought to elucidate the integration of control and mirroring processes, examining possible routes for the contextual gating of visuomotor information during intentional and incidental (automatic) mirroring. For the current task, pitting intentional against incidental SR mapping, higher order response-inhibition and goal-directed processes are required to explain the observed interaction effect within a parietal-occipital region of the action observation network. In particular, we sought to disambiguate amongst competing mechanisms for this congruence-by-intentionality effect, given the representations of SR mismatch in the anterior insula. We found that the insula played a pivotal role, gating connectivity between the SMA and MOG nodes, so that the visual representation of mirror-matched actions was dampened when it was either task-irrelevant (pre-defined responses incidental to observed actions) or counter-productive to explicit goals (intentional counter-imitation). These findings expand our understanding of the neural basis of one of the most elemental of human behaviours – mirroring – and its placement within the context of cognitive control and variable task demands. They also provide a benchmark for studies of disturbances to mirroring as implicated in a myriad of disorders, including schizophrenia (Lee et al., 2014), autism (Perkins et al., 2010) and echopraxia (Ganos et al., 2012).

As a whole, the mirror neuron system is posited to encompass both higher-order understanding, such as goals and intentions underlying actions (Kilner et al., 2007a; Oztop et al., 2013; Sinigaglia, 2013), and lower-level, more automatic sensorimotor integration of the kinematics and visual perception of actions (D’Ausilio et al., 2015; Kilner et al., 2003). This low-level of sensorimotor integration has been established in the ‘automatic imitation’ behavioural effect during incidental action observation (Heyes, 2011). Beyond this more automatic mapping, the question remains of how top-down influences can guide the way mirror representations are engaged during intentional response preparation. Previous evidence of controlled mirroring includes preparatory suppression of corticospinal excitability, in which the neuromuscular mapping of observed actions is dampened, both in advance of executing a non-imitative action (Cross and Iacoboni, 2014b; 2014a) and viewing action stimuli passively after having previously been instructed to prepare non-imitative responses (Bardi et al., 2015). These measures of response suppression for non-mirrored actions show that the tendency for automatic imitation can be bent to task demands and intentions. Here, we have offered a model for how this controlled mirroring can occur, through the engagement of executive control networks. By directly comparing pro-active control processes during intentional counter-imitation to reactive control during incidental counter-imitation, as well as equivalent imitative contexts, our paradigm uniquely targeted the task-dependent control of this imitative mirroring tendency. The modulation of effective connectivity between the SMA and MOG, by the anterior insula, accounted for the cortical activity distinctly evoked by intentional rather than incidental SR mapping. This integration of action observation and executive control networks explains how the very human capacity for mirroring other agents can be deployed adaptively to optimise our responses to ever changing demands.

Task-relevance was intended to target executive control and higher-order representations of SR relationships. Such controlled mirroring is crucial to real-world interactions, in which context offers necessary cues to control our more automatic tendency to ‘mirror’ a partner’s action. Even if mirroring were *solely* congruent (which is unlikely), it is necessary that humans are capable of inhibiting mirroring for normal social functioning. Whilst some degree of mimicry has been shown to increase social cohesion and rapport (Hamilton, 2013; Chartrand and Bargh, 1999; Lakin & Chartrand, 2003) “mimicking the passive unconscious behaviours of another person is tantamount to mocking” (p. 338, Lakin & Chartrand, 2003). Indeed, excessive or exaggerated imitation may indicate a disorder of response inhibition, for example extremely disordered imitation indicative of echopraxia. This rare condition occurs in patients with frontal lobe damage who repeat all observed actions, with such movements being involuntary, reflexive and automatic (Ganos et al., 2012; Lhermitte et al., 1986). Our model concurs with the anterior frontal lobe being required for intentional suppression of mirroring during passive action observation (Bien et al., 2009), the opposite of the compulsive imitation observed in echopraxia. Moreover, Bien and colleagues (Bien et al., 2009) have described a general pattern of response inhibition relating to information flow from mid-frontal and insula areas via premotor to parietal and temporal-occipital cortices, i.e. from frontal control back through fronto-parietal networks, along an anterior (motor) to posterior (visual) pathway. We also highlighted a hierarchy for effectivity connectivity between the anterior insula over the SMA and influencing communication with the MOG.

There are several caveats of our study. Crucially, we have adopted one particular use of DCM, integrating specific hypotheses into a relatively restrictive model space. Indeed, our models were purposely constrained to as few regions of interest as possible to explain the BOLD responses evoked by our task, yet still capturing visual, motor, mirror, and executive control functions. Nevertheless, as with any study, other hypotheses could have been entertained. For example, we did not include other regions of the control system in our model network, such as the inferior frontal gyrus pars operculum (IFGpo). Past findings of the IFGpo being engaged during controlled mirroring have positioned this as the frontal node of the mirror neuron system providing a connection with executive control networks (Cross et al., 2013). This previous study focused on effective connectivity within the frontal control networks (ACC, mPFC, aINS), to this frontal mirror region (IFGpo), and produced evidence of SR conflict (mismatch) modulating the anterior insula connection with the IFGpo (Cross et al., 2013). Our streamlined DCM designs do not preclude the effective connections of this previous model (Cross et al., 2013). Indeed, given that the IFGpo is itself implicated in frontal control networks (Cai et al., 2014b; Dosenbach et al., 2008), with co-activation of the anterior insula and IFGpo being a consistent finding in response-inhibition (Dosenbach et al., 2008; 2007), we acknowledge that the IFG may also contribute to control processes as modelled here. Nonetheless, we have clearly shown a crucial role of the anterior insula in this modulatory process. Specifically, the anterior insula influenced the integration of motor preparation from the SMA with the visual representations of actions in the MOG. This agrees with previous work which marked the anterior insula as integral for detecting behaviourally salient stimuli and successful response-inhibition within the cingulo-opercula control network (Cai et al., 2014b). Thus, we have disentangled specific detail of where top-down control is exerted amongst the interrelated processes of action observation, mirroring and motor planning.

Another point of note is that our models did not explicitly include a node within a canonical mirror area. Having taken a data-driven approach for defining our regions of interest with the explicit aim of describing the interaction effect observed, the ‘nodes’ of our model were restricted to the local maxima detected by the GLM analysis. For the parietal-occipital cluster linked to the key congruence-intentionality interaction, this data-driven approach placed the region of interest in the occipital portion and not the parietal portion. Nevertheless, dynamic causal modelling does not assume that all effects are described by the network nodes, rather it allows for other non-specified intermediate regions to be encompassed by those nodes and connections that are explicitly defined (Zeidman et al., 2019). Moreover, the importance of regions of a wider action observation network including temporal-occipital areas have often been overlooked by research focused on frontoparietal regions (Lingnau and Downing, 2015). Thus, our present findings explain the modulation of action observation processing beyond the classical frontoparietal mirror network. This suggest that these regions of the ‘action-observation network’ play an important role in how sensorimotor representations are integrated during imitation and counter imitation.

The human capacity for mirroring observed actions is integrated with higher-order cognitive processes in a more complex and nuanced way than a direct one-to-one matching of stimulus and response movements. Our behavioural paradigm targeted the task-relevance of mirror-matched representations of observed actions for planning self-made movements. We have highlighted one crucial aspect of this integrative process, namely the task-relevance or intentionality of performing imitative actions. The engagement of executive control processes appears to gate the interactions between the SMA and MOG through nonlinear modulation exerted by the insula cortex. These findings point to several avenues for future work. For instance, it would be interesting to explore the temporal dynamics of this hierarchical gating of the representations of observed actions through use of imaging methods with higher temporal resolution (combined EEG-fMRI or MEG). Subtle differences in the timing and the wave-forms of evoked responses in these regions could allow our findings to be positioned within the broader theoretical framework of predictive coding (Kilner et al., 2007a; 2007b). An alternative extension would be to reframe mirroring within a hierarchy of other domain-general processes. For example, associative learning processes are thought to underlie the formation of mirror associations (Heyes et al., 2005), yet the confluence of task-relevance with sensorimotor experience is yet to be examined. This would assess whether the hierarchical role of the anterior insula in context-dependent motor mirroring, as shown here, may form a crucial circuit for more general processes that could similarly modulate the engagement of other learnt sensorimotor associations and not just imitative actions.

## Acknowledgements

Australian Research Council Special Research Initiative (SR12030015) Science of Learning Research Centre funding supported this study, and while conducting this work the first author received an Australian Postgraduate Award. This work was conducted as part of the first author’s PhD project and she gladly acknowledges the support and guidance of her supervisors, academic review panel and the wider cog-neuro community at UQ. Immense gratitude is owed to Mr Aiman Al Najjar (Human Imaging Manager at the Centre for Advanced Imaging, The University of Queensland), Dr David Lloyd (The Queensland Brain Institute) and Dr. Chase Sherwell (School of Psychology, The University of Queensland) for their technical expertise and unwavering encouragement.

## Supplementary Information

### S1. Methods - Further detail on DCM specification

A DCM is a collection of nodes (regions) and the directed connections between them. For task fMRI, VOIs are selected as network nodes based on the task effects in the data (see Methods). DCMs are then specified by 4 matrices which are the terms in the neural state-equation (Friston et al, 2003; Marreiros, Stephan, & Friston, 2010; Stephan et al. 2010) If *n* is the number of nodes and *i* the number of inputs then these matrices are,

- A-matrix: an [*n* x *n]* matrix of intrinsic connections (edges between nodes) for inter- and intra-node communication, can be framed as bidirectional/unidirectional or feed-back/- forward and lateral connections.
- B-matrix: an [*n* x *n* x *i*] matrix of task-related modulator effects, induced context-dependent changes in connectivity; bilinear term state equation. For example: a second-order interaction between the driving inputs (task conditions) and activity in a source node related to a response in a target node. See Results Section 3.1.
- C-matrix: an [*n* x *i*] matrix for the driving inputs based on the known perturbations of the experimental paradigm, e.g. stimulus properties/ task conditions.
- D-matrix: an [*n* x *n* x *n*] matrix for nonlinear modulations from one node onto the connection between 2 other nodes. Note these D-matrices are only specified for nonlinear DCMs and not included in bilinear DCMs. See Results Section 3.1.

For the present study, the matrices across all 48 DCM designs were specified using custom MATLAB batchscripts to call the DCM functions from the SPM12 toolbox.

#### S1.1 Node selection from GLM and intrinsic connections

Four regions of interest were selected as nodes included in all DCM hypotheses: The node for V5/MT was defined by the peak signal-change (p<0.05 FWE corrected) across all task conditions over baseline within the AAL defined left middle temporal gyrus (MNI coordinates: [−44 −66, 12]; note the same peak was also identified when WFU Pickatlas (Maldjian et al., 2003) toolbox was used to define the ROI with left MT mask). This region was chosen as it is an area within early visual cortex sensitive to visual motion and it responded strongly to all four task conditions (see Results Section 3 Fig. 3.B for beta-estimate plots). It was hence modelled as receiving driving inputs related to visual movement stimuli across all conditions. The MOG peak [−28, −66, 26] was highlighted by an interaction effect driven by the factorial combination intentional imitation. SMA is known to be critical for motor planning, and a peak [−4, 14, 48] identified for the main effect of action-preparation was used for the third VOI. Finally, the VOI within the left anterior insula cortex [−30, 20, 2] was based on the main effect of SRC and interpreted to reflect conflict-detection, response-inhibition and goal maintenance processing within the cingulo-opercula network for cognitive control (Dosenbach et al., 2008; Nelson et al., 2010).

#### S1.2 Specification of intrinsic network edges (A-matrix)

Across all models, two areas were positioned as ‘fully’ connected with all other network nodes: the MOG and the insula. For example, in our base design for connections (DCM A-matrix), the MOG was bi-directionally linked to the insula node and to both the SMA and V5MT, while these latter two regions were themselves not directly connected.

This design was informed by literature on the functional and structural connectivity of these particular areas. The peak coordinate centred on for the **MOG node** of our models was located near the intraparietal sulcus and angular gyrus of the IPL. This region is a confluence of temporo-parieto-occipital cortices, where subcortical white matter tracts form a crucial hub of intra- and inter-cortical structural connectivity providing high-level multimodal integration (De Benedictis et al., 2014). Relevant to action observation processes, functional connectivity between the insula, parietal, and occipital cortices are integrated with a general posterior to anterior organisation. This organisation has been argued to allow the transformation of stimuli from IPL-occipital connections through to anterior IPL connections with the prefrontal cortex and insula cortex (Uddin et al., 2010). Given this prior information, our DCM hypotheses framed the MOG node as being “fully” connected within our models.

Regarding the **anterior insula cortex**, this area is involved in a very wide range of tasks (Craig, 2010). The insula cortex has been firmly posited as an integral hub in the salience network, via strong coupling with the anterior cingulate cortex to facilitate rapid links to the motor system and assist appropriate responses to salient stimuli (Menon and Uddin, 2010). This fits well with the insula cortex being inclused within the cingulo-opercula control network (Dosenbach et al., 2008). This cingulo-opercula network is comprised of the anterior insula cortex, the anterior prefrontal cortex, frontal operculum, dorsal anterior cingulate cortex and thalamus; and is argued to function in counter-part to a fronto-parietal network, together framed as dual-networks for cognitive control (Cai et al., 2014a; Dosenbach et al., 2008). Structurally, the anterior insula connects with posterior insula as well as frontal, parietal, and temporal lobes (Faillenot et al., 2017), and bilaterally with the SMA (Cauda et al. 2011). These details informed our A-matrix design for the insula node being modelled as connected with all the other regions in our model network.

**The SMA** has been reliably related to higher order motor planning and preparation for voluntary, intentional movements (Cunnington et al., 2005). Coupled with our task effects on BOLD signal changes highlighting involvement of the SMA for intentional compared with incidental action contexts, we viewed this as representing higher order planning for task-relevant SR pairs. Thus, we hypothesised this node as effectively connected with higher order regions of our network, the insula and MOG, but not with V5/MT (low-level perception of visual motion).

#### S1.3 Specification of driving inputs (C-matrix)

Our inclusion of area V5 was motivated as a low-level visual representation of movements, (ffytche et al., 1995; Zeki, 2016), as a bottom-up sensory signal for the observed action common to all task conditions. Thus, driving inputs to this node were ubiquitous to all hypothesised networks. Another potential driving input considered was early motor preparation which was modelled as a driving input to the SMA in a family of dual-input designs (Results Fig.3A).

### S2 Results – Peak statistics for whole-brain GLM analysis

**Table S1.**
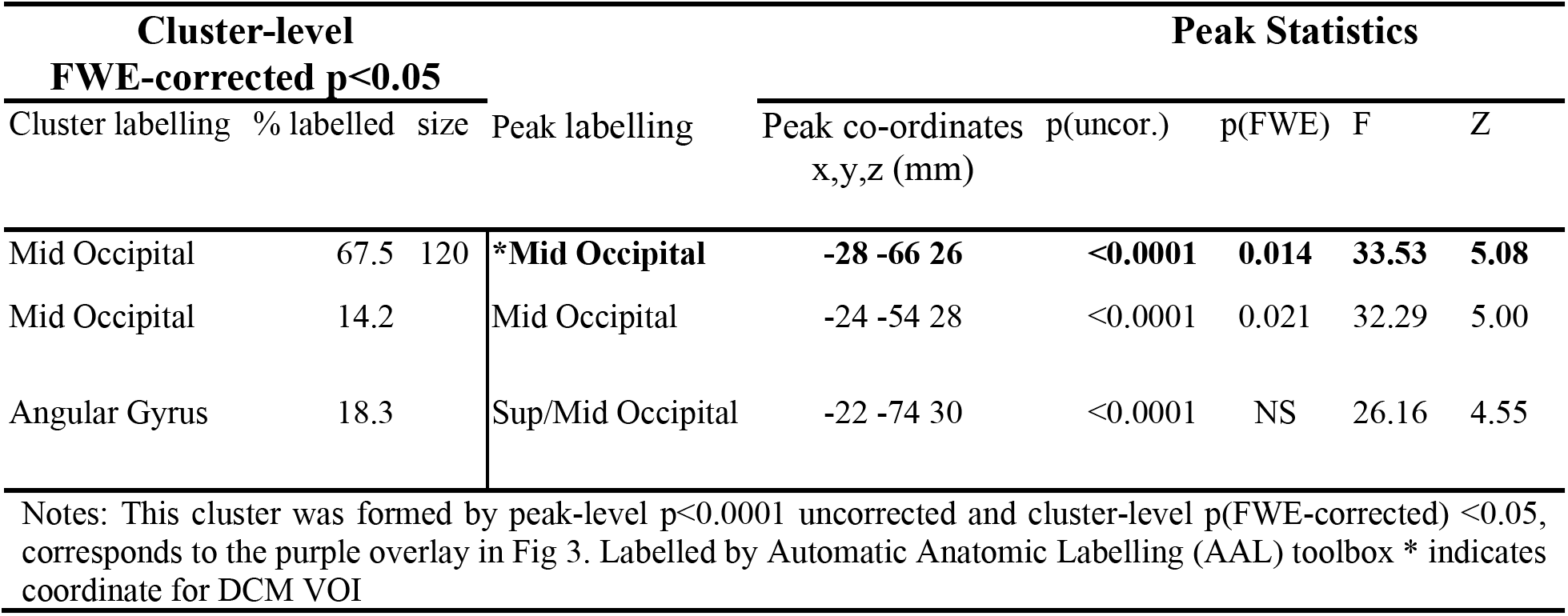
Summary of peak statistics interaction effects (task-relevance x congruence) on task-related BOLD responses. Cluster-level FWE-correction p<0.05.

| Cluster-level<br>FWE-corrected $p < 0.05$ | | | Peak Statistics | | | | | |
| --- | --- | --- | --- | --- | --- | --- | --- | --- |
| Cluster labelling | % labelled | size | Peak labelling | Peak co-ordinates<br>x,y,z (mm) | p(uncor.) | p(FWE) | F | Z |
| Mid Occipital | 67.5 | 120 | **\*Mid Occipital** | **-28 -66 26** | **<0.0001** | **0.014** | **33.53** | **5.08** |
| Mid Occipital | 14.2 |  | Mid Occipital | -24 -54 28 | <0.0001 | 0.021 | 32.29 | 5.00 |
| Angular Gyrus | 18.3 |  | Sup/Mid Occipital | -22 -74 30 | <0.0001 | NS | 26.16 | 4.55 |

Notes: This cluster was formed by peak-level $p < 0.0001$ uncorrected and cluster-level $p(\text{FWE-corrected}) < 0.05$ , corresponds to the purple overlay in Fig 3. Labelled by Automatic Anatomic Labelling (AAL) toolbox \* indicates coordinate for DCM VOI

**Table S2.**
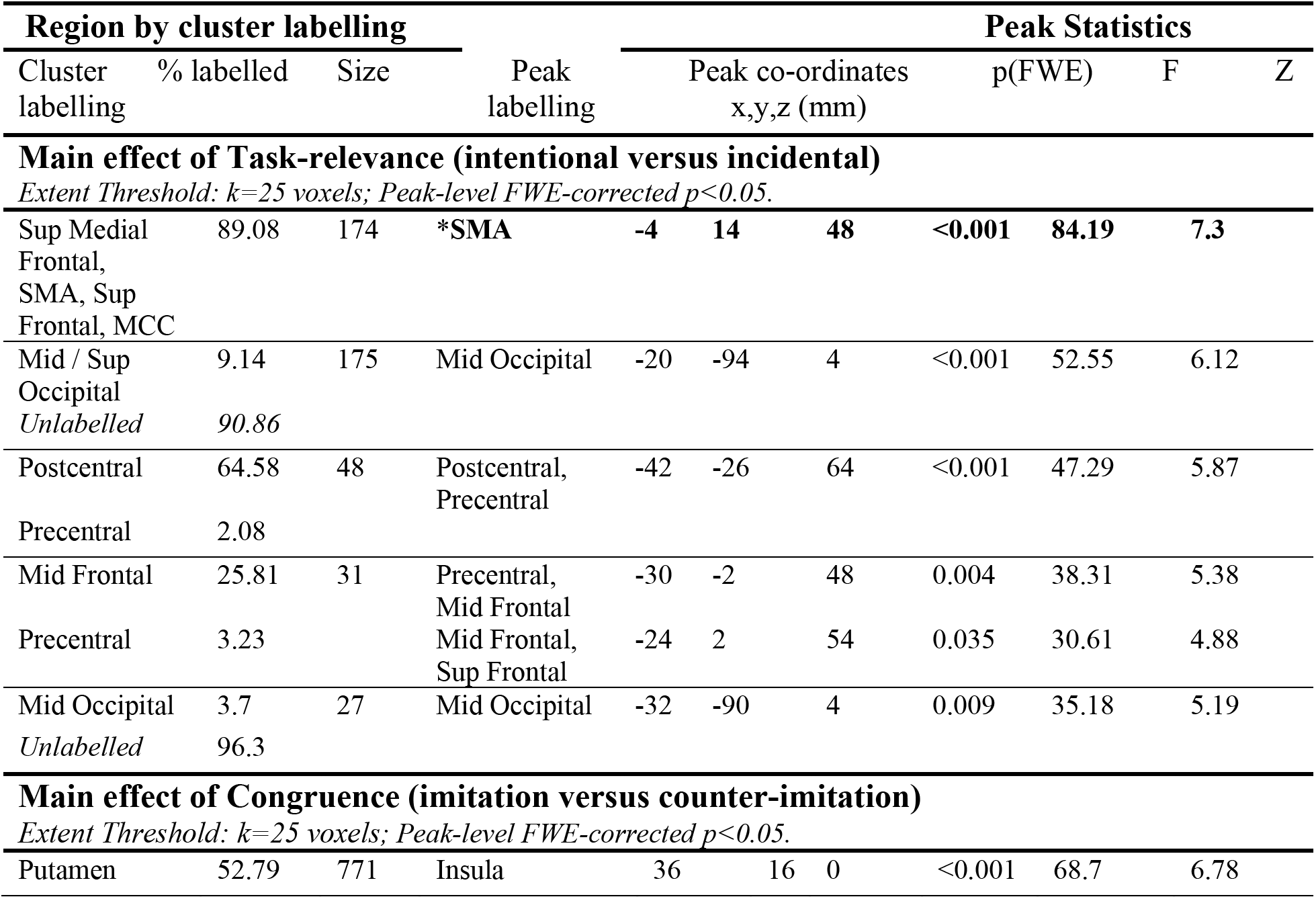
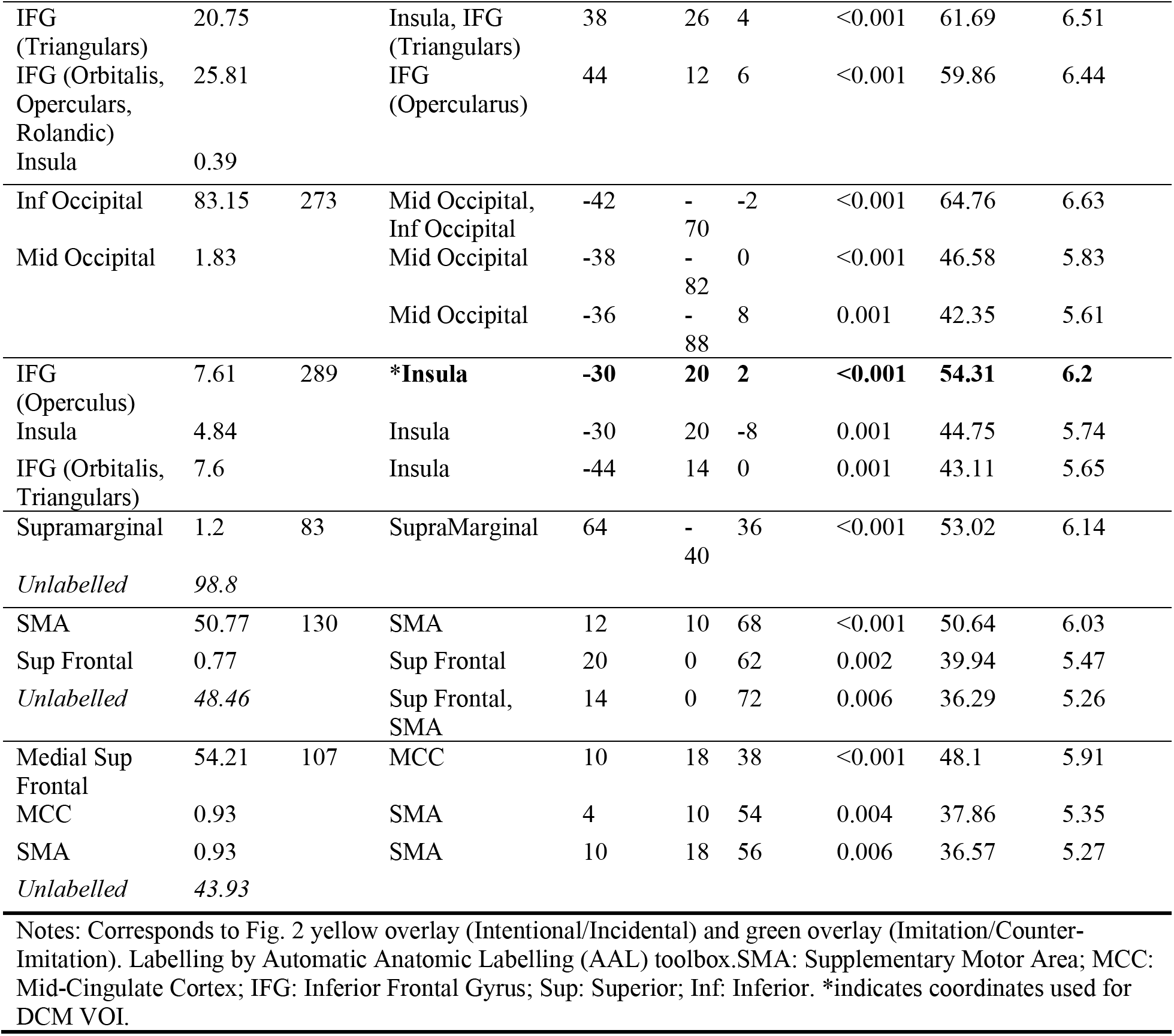
Summary of peak statistics for main effects from whole-brain analysis of task-related BOLD responses. Peak-level FWE-correction p<0.05.

## References

Bardi, L., Bundt, C., Notebaert, W., Brass, M., 2015. Eliminating mirror responses by instructions. Cortex 70, 128–136. doi:10.1016/j.cortex.2015.04.018

Bien, N., Roebroeck, A., Goebel, R., Sack, A.T., 2009. The Brain’s Intention to Imitate: The Neurobiology of Intentional versus Automatic Imitation. Cerebral Cortex 19, 2338–2351. doi:10.1093/cercor/bhn251

Brass, M., Bekkering, H., Wohlschläger, A., Prinz, W., 2000. Compatibility between Observed and Executed Finger Movements: Comparing Symbolic, Spatial, and Imitative Cues. Brain and Cognition 44, 124–143. doi:10.1006/brcg.2000.1225

Brass, M., Derrfuss, J., Cramon, von, D.Y., 2005. The inhibition of imitative and overlearned responses: a functional double dissociation. Neuropsychologia 43, 89–98. doi:10.1016/j.neuropsychologia.2004.06.018

Brass, M., Ruby, P., Spengler, S., 2009. Inhibition of imitative behaviour and social cognition. Philosophical Transactions of the Royal Society of London B: Biological Sciences 364, 2359–2367. doi:10.1098/rstb.2009.0066

Brass, M., Zysset, S., Cramon, von, D.Y., 2001. The inhibition of imitative response tendencies. NeuroImage 14, 1416–1423. doi:10.1006/nimg.2001.0944

Cai, W., Ryali, S., Chen, T., Li, C.-S.R., Menon, V., 2014a. Dissociable Roles of Right Inferior Frontal Cortex and Anterior Insula in Inhibitory Control: Evidence from Intrinsic and Task-Related Functional Parcellation, Connectivity, and Response Profile Analyses across Multiple Datasets. Journal of Neuroscience 34, 14652–14667. doi:10.1523/JNEUROSCI.3048-14.2014

Cai, W., Ryali, S., Chen, T., Li, C.S.R., Menon, V., 2014b. Dissociable Roles of Right Inferior Frontal Cortex and Anterior Insula in Inhibitory Control: Evidence from Intrinsic and Task-Related Functional Parcellation, Connectivity, and Response Profile Analyses across Multiple Datasets. Journal of Neuroscience 34, 14652–14667. doi:10.1523/JNEUROSCI.3048-14.2014

Campbell, M.E.J., Cunnington, R., 2017. More than an imitation game: Top-down modulation of the human mirror system. Neuroscience and Biobehavioral Reviews 75, 195–202. doi:10.1016/j.neubiorev.2017.01.035

Campbell, M.E.J., Mehrkanoon, S., Cunnington, R., 2018. Intentionally not imitating: Insula cortex engaged for top-down control of action mirroring. Neuropsychologia 111, 241–251. doi:10.1016/j.neuropsychologia.2018.01.037

Caspers, S., Zilles, K., Laird, A.R., Eickhoff, S.B., 2010. ALE meta-analysis of action observation and imitation in the human brain. NeuroImage 50, 1148–1167. doi:10.1016/j.neuroimage.2009.12.112

Catmur, C., 2016. Automatic imitation? Imitative compatibility affects responses at high perceptual load. J Exp Psychol Hum Percept Perform 42, 530–539. doi:10.1037/xhp0000166

Cauda, F., D’Agata, F., Sacco, K., Duca, S., Geminiani, G., & Vercelli, A. (2011). Functional connectivity of the insula in the resting brain. NeuroImage, 55(1), 8–23. doi:10.1016/j.neuroimage.2010.11.049

Chartrand, T.L., Bargh, J.A., 1999. The chameleon effect: The perception-behavior link and social interaction. Journal of Personality and Social Psychology 76, 893–910. doi:10.1037/0022-3514.76.6.893

Choi, H.-J., Zilles, K., Mohlberg, H., Schleicher, A., Fink, G.R., Armstrong, E., Amunts, K., 2006. Cytoarchitectonic identification and probabilistic mapping of two distinct areas within the anterior ventral bank of the human intraparietal sulcus. J. Comp. Neurol. 495, 53–69. doi:10.1002/cne.20849

Craig, A.D., 2010. Once an island, now the focus of attention. Brain Structure & Function 214, 395–396. doi:10.1007/s00429-010-0270-0

Cross, K.A., Iacoboni, M., 2014a. Neural systems for preparatory control of imitation. Philos. Trans. R. Soc. Lond., B, Biol. Sci. 369, 20130176–20130176. doi:10.1016/j.bbr.2008.08.028

Cross, K.A., Iacoboni, M., 2014b. To imitate or not: Avoiding imitation involves preparatory inhibition of motor resonance. NeuroImage 91, 288–236.

Cross, K.A., Torrisi, S., Reynolds Losin, E.A., Iacoboni, M., 2013. Controlling automatic imitative tendencies: Interactions between mirror neuron and cognitive control systems. NeuroImage 83, 493–504. doi:10.1016/j.neuroimage.2013.06.060

Cunnington, R., Windischberger, C., Moser, E., 2005. Premovement activity of the pre-supplementary motor area and the readiness for action: Studies of time-resolved event-related functional MRI. Human Movement Science 24, 644–656. doi:10.1016/j.humov.2005.10.001

D’Ausilio, A., Bartoli, E., Maffongelli, L., 2015. Grasping synergies: a motor-control approach to the mirror neuron mechanism. Physics of Life Reviews 12, 91–103. doi:10.1016/j.plrev.2014.11.002

De Benedictis, A., Duffau, H., Paradiso, B., Grandi, E., Balbi, S., Granieri, E., Colarusso, E., Chioffi, F., Marras, C.E., Sarubbo, S., 2014. Anatomo-functional study of the temporo-parieto-occipital region: dissection, tractographic and brain mapping evidence from a neurosurgical perspective. Journal of Anatomy 225, 132–151. doi:10.1111/joa.12204

Dosenbach, N.U.F., Fair, D.A., Cohen, A.L., Schlaggar, B.L., Petersen, S.E., 2008. A dual-networks architecture of top-down control. Trends in Cognitive Sciences 12, 99–105. doi:10.1016/j.tics.2008.01.001

Dosenbach, N.U.F., Fair, D.A., Miezin, F.M., Cohen, A.L., Wenger, K.K., Dosenbach, R.A.T., Fox, M.D., Snyder, A.Z., Vincent, J.L., Raichle, M.E., Schlaggar, B.L., Petersen, S.E., 2007. Distinct brain networks for adaptive and stable task control in humans. Proceedings of the National Academy of Sciences 104, 11073–11078. doi:10.1073/pnas.0704320104

Faillenot, I., Heckemann, R.A., Frot, M., Hammers, A., 2017. Macroanatomy and 3D probabilistic atlas of the human insula. NeuroImage 150, 88–98. doi:10.1016/j.neuroimage.2017.01.073

Ferrari, P.F., Rizzolatti, G., 2014. Mirror neuron research: the past and the future. Philosophical Transactions of the Royal Society of London B: Biological Sciences 369 (1644), 1–4. doi:10.1098/rstb.2013.0169.

ffytche, D.H., Guy, C.N., Zeki, S., 1995. The parallel visual motion inputs into areas V1 and V5 of human cerebral cortex. Brain 118 (Pt 6), 1375–1394. doi:10.1093/brain/118.6.1375

Friston, K.J., Harrison, L., Penny, W., 2003a. Dynamic causal modelling. NeuroImage 19, 1273–1302. doi:10.1016/S1053-8119(03)00202-7

Ganos, C., Ogrzal, T., Schnitzler, A., Münchau, A., 2012. The pathophysiology of echopraxia/echolalia: relevance to Gilles de la Tourette syndrome. Mov. Disord. 27, 1222–1229. doi:10.1002/mds.25103

Hamilton, A.F.deC., (2013). The mirror neuron system contributes to social responding. Cortex 49(10), 2957–2959. doi:10.1016/j.cortex.2013.08.012

Heyes, C., 2011. Automatic imitation. Psychological Bulletin 137, 463. doi:10.1037/a0022288

Heyes, C., Bird, G., Johnson, H., Haggard, P., 2005. Experience modulates automatic imitation. Cognitive Brain Research 22, 233–240. doi:10.1016/j.cogbrainres.2004.09.009

Kahan, J., Foltynie, T., 2013. Understanding DCM: Ten simple rules for the clinician. NeuroImage 83, 542–549. doi:10.1016/j.neuroimage.2013.07.008

Kilner, J.M., Friston, K.J., Frith, C.D., 2007a. The mirror-neuron system: a Bayesian perspective. Neuroreport 18, 619–623. doi:10.1097/WNR.0b013e3281139ed0

Kilner, J.M., Friston, K.J., Frith, C.D., 2007b. Predictive coding: an account of the mirror neuron system. Cogn Process 8, 159–166. doi:10.1007/s10339-007-0170-2

Kilner, J.M., Paulignan, Y., Blakemore, S.J., 2003. An interference effect of observed biological movement on action. Current Biology 13, 522–525. doi:10.1016/S0960-9822(03)00165-9

Lakin, J.L., Chartrand, T.L. (2003), Using nonconscious behavioral mimicry to create affiliation and rapport. Psychol. Sci. 14(4), 334–339. Doi:10.1111/1467-9280.14481

Lee, J.S., Chun, J.W., Yoon, S.Y., Park, H.-J., Kim, J.-J., 2014. Involvement of the mirror neuron system in blunted affect in schizophrenia. Schizophr Res 152, 268–274. doi:10.1016/j.schres.2013.10.043

Lhermitte, F., Pillon, B., Serdaru, M., 1986. Human autonomy and the frontal lobes. Part I: Imitation and utilization behavior: A neuropsychological study of 75 patients. Annals of Neurology 19, 326–334. doi:10.1002/ana.410190404

Lingnau, A., Downing, P.E., 2015. The lateral occipitotemporal cortex in action. Trends in Cognitive Sciences 19, 268–277. doi:10.1016/j.tics.2015.03.006

Maldjian, J.A., Laurienti, P.J., Burdette, J.B., Kraft, R.A., 2003. An Automated Method for Neuroanatomic and Cytoarchitectonic Atlas-based Interrogation of fMRI Data Sets. NeuroImage 1233–1239. doi:10.1016/S1053-8119(03)00169-1

Marreiros, A.C., Stephan, K.E., Friston, K.J., 2010. Dynamic causal modeling. Scholarpedia 5, 9568. doi:10.4249/scholarpedia.9568

Mehrkanoon, S., Breakspear, M., Boonstra, T.W., 2014. The reorganization of corticomuscular coherence during a transition between sensorimotor states. NeuroImage 100, 693–702. doi:10.1016/j.neuroimage.2014.06.050

Menon, V., Uddin, L.Q., 2010. Saliency, switching, attention and control: a network model of insula function. Brain Struct Funct 214, 655–667. doi:10.1007/s00429-010-0262-0

Molenberghs, P., Cunnington, R., Mattingley, J.B., 2012. Brain regions with mirror properties: A meta-analysis of 125 human fMRI studies. Neuroscience Biobehav Rev 36, 341–349. doi:10.1016/j.neubiorev.2011.07.004

Molenberghs, P., Cunnington, R., Mattingley, J.B., 2009. Is the mirror neuron system involved in imitation? A short review and meta-analysis. Neuroscience Biobehav Rev 33, 975–980. doi:10.1016/j.neubiorev.2009.03.010

Nelson, S.M., Dosenbach, N.U.F., Cohen, A.L., Wheeler, M.E., Schlaggar, B.L., Petersen, S.E., 2010. Role of the anterior insula in task-level control and focal attention. Brain Structure & Function 214, 669–680. doi:10.1007/s00429-010-0260-2

Oztop, E., Kawato, M., Arbib, M.A., 2013. Mirror neurons: Functions, mechanisms and models. Neuroscience letters 540, 43–55. doi:10.1016/j.neulet.2012.10.005

Penny, W.D., 2012. Comparing dynamic causal models using AIC, BIC and free energy. NeuroImage 59, 319–330. doi:10.1016/j.neuroimage.2011.07.039

Perkins, T., Stokes, M., McGillivray, J., Bittar, R., 2010. Mirror neuron dysfunction in autism spectrum disorders. Journal of Clinical Neuroscience 17, 1239–1243. doi:10.1016/j.jocn.2010.01.026

Prinz, W., 1997. Perception and action planning. Eur. J. Cogn. Psychol. 9, 129–154, http://dx.doi.org/10.1080/713752551.

Rizzolatti, G., Sinigaglia, C., 2010. The functional role of the parieto-frontal mirror circuit: interpretations and misinterpretations. Nat. Rev. Neurosci. 11, 264–274. doi:doi:10.1038/nrn2805

Sinigaglia, C., 2013. What type of action understanding is subserved by mirror neurons? Neuroscience letters 540, 59–61. doi:10.1016/j.neulet.2012.10.016

Stephan, K.E., Friston, K.J., 2010. Analyzing effective connectivity with functional magnetic resonance imaging. WIREs Cogn Sci 1, 446–459. doi:10.1002/wcs.58

Stephan, K.E., Penny, W.D., Moran, R.J., Ouden, den, H.E.M., Daunizeau, J., Friston, K.J., 2010. Ten simple rules for dynamic causal modeling. NeuroImage 49, 3099–3109. doi:10.1016/j.neuroimage.2009.11.015

Stephan, K.E., Kasper, L., Harrison, L.M., Daunizeau, J., Ouden, den, H.E.M., Breakspear, M., Friston, K.J., 2008. Nonlinear dynamic causal models for fMRI. NeuroImage 42, 649–662. doi:10.1016/j.neuroimage.2008.04.262

Stephan, K.E., Penny, W.D., Daunizeau, J., Moran, R.J., Friston, K.J., 2009. Bayesian model selection for group studies. NeuroImage 46, 1004–1017. doi:10.1016/j.neuroimage.2009.03.025

Tzourio-Mazoyer, N., Landeau, B., Papathanassiou, D., Crivello, F., Etard, O., Delcroix, N., Mazoyer, B., Joliot, M., 2002. Automated anatomical labeling of activations in SPM using a macroscopic anatomical parcellation of the MNI MRI single-subject brain. NeuroImage 15, 273–289. doi:10.1006/nimg.2001.0978

Uddin, L.Q., Supekar, K., Amin, H., Rykhlevskaia, E., Nguyen, D.A., Greicius, M.D., Menon, V., 2010. Dissociable connectivity within human angular gyrus and intraparietal sulcus: Evidence from functional and structural connectivity. Cerebral Cortex 20, 2636–2646. doi:10.1093/cercor/bhq011

Zeidman, P., Jafarian, A., Corbin, N., Seghier, M. L., Razi, A., Price, C. J., & Friston, K. J. (2019). A guide to group effective connectivity analysis, part 1: First level analysis with DCM for fMRI. NeuroImage 200, 174–190. doi:10.1016/j.neuroimage.2019.06.031.

Zeki, S., 2016. Parallel Processing, Asynchronous Perception, and a Distributed System of Consciousness in Vision. The Neuroscientist. doi:10.1177/107385849800400518

Zwickel, J., Prinz, W., 2012. Assimilation and contrast: the two sides of specific interference between action and perception. Psychol. Res. 76, 171–182, http://dx.doi.org/10.1007/s00426-011-0338-3.

## Supplementary Information References

Dosenbach, N.U.F., Fair, D.A., Cohen, A.L., Schlaggar, B.L., Petersen, S.E., 2008. A dual-networks architecture of top-down control. Trends in Cognitive Sciences 12, 99–105. doi:10.1016/j.tics.2008.01.001

Friston, K.J., Harrison, L., Penny, W., 2003b. Dynamic causal modelling. NeuroImage 19, 1273–1302. doi:10.1016/S1053-8119(03)00202-7

